# State diagrams of type I collagen for the rational design of biomimetic materials

**DOI:** 10.1101/2024.05.17.594626

**Authors:** Isabelle Martinier, Sylvain Deville, Gervaise Mosser, Léa Trichet, Patrick Davidson, Francisco M. Fernandes

## Abstract

Ideally, designing tissue engineering grafts and 3D cell culture materials should rely on mimicking the architecture and composition of the extracellular matrix, which is predominantly comprised of type I collagen. However, while collagen molecules are assembled into fibrils by cells in vivo, well-organized fibrils rarely form spontaneously in vitro. Indeed, the physico-chemical conditions for fibrillogenesis are still poorly understood and their influence on the formation and properties of fibrillar biomimetic materials remains elusive. Here, we establish state diagrams for type I collagen over an unprecedented range of concentration and temperature, showing the collagen denaturation limits, the emergence of fibrils in acidic conditions, and a new regime of collagen molecule/fibril coexistence. We also show how the state diagrams can be used to understand the formation of biomimetic materials by classical methods, as illustrated here by collagen freeze-casting. Therefore, these state diagrams will help to optimize the production of collagen-based biomimetic materials.

## Introduction

*In vivo*, the supramolecular assembly of type I collagen molecules into fibrils underpins the formation of the Extracellular Matrix (ECM), a central aspect of tissue morphogenesis and remodeling. This process occurs in a narrow range of physicochemical conditions (temperature, concentration, and ionic strength) that are defined by the physiology of the cells and by the composition of the extracellular environment. We will henceforth refer to this set of conditions as the *physiological physicochemical range*. The current paradigm in collagen-based biomimetic materials states that reproducing the same physicochemical conditions *in vitro* as *in vivo*, is the key to mimicking the architecture—and functions—of biological tissues^1^. This approach is gaining traction as it promises to tailor materials for applications such as 3D cell culture models^2^ or prosthetic grafts^3,4^.

The state of the art is rich in reports describing biomaterials built with collagen as the central building block to mimic the ECM. *In vitro*, fibrillogenesis, the transformation of collagen solutions into fibrillar gels, is an entropy-driven process that occurs above 1 nM through a change in pH or ionic strength^5^. This step triggers the formation of fibrils, which are the building blocks recognized by cells and that contribute to the mechanical properties of tissues. In fibrils, hydrated type I collagen molecules are arranged in a regular staggered parallel pattern, with an axial periodicity of 67 nm known as the D-banding period^6^. Preserving this subtle self-assembly in material fabrication techniques requires a deep understanding of its mechanisms. Molecular and supramolecular order is determined by weak interactions between collagen molecules—electrostatic, van der Waals, hydrophobic, and hydrogen-bonds^7^—and follows a liquid-crystal-like behavior, as first described by Y. Bouligand^8^. Since then, the fluid phases of collagen (isotropic, nematic, cholesteric), which occur in the vicinity of the physiological physicochemical range, have been explored for a wide range of collagen concentrations^9^, acid types and their respective concentrations^10,11^, using microscopy tools (polarized light optical microscopy, PLOM, and transmission electron microscopy, TEM^9,12–14^).

In parallel, the development of biomimetic collagenous materials has increasingly been coupled with materials processing techniques such as ice-templating^15–21^, electrospinning^22,23^, spray-drying^24,25^, 3D bioprinting^26–30^, electrochemically aligned collagen (ELAC)^31^ or cell-mediated tissue engineering^32,33^. Many of these reports refer to concentrations below those of native tissues and processing temperatures above the denaturation temperature of collagen molecules, resulting in particularly expensive gelatin-based materials as discussed elsewhere^34^. The native structures of collagen are thus rarely achieved by most of materials engineering strategies, due to a large gap between the bioengineering community and the basic research community focused on biological self-assembly mechanisms (structural biologists, biochemists and biophysicists). As a result, there is a significant lack of guidance on how to preserve the physiological physicochemical range in harsh processes aimed at mimicking biological tissues^35^. Incorporating the conditions that ensure collagen self-assembly into harsh processing strategies that rely on drastic temperature and concentration changes (e.g. evaporation or phase separation induced by ice growth) remains a major challenge. Simply put, the underlying question can be formulated as follows:

### How can the weak interactions that govern the supramolecular assembly of collagen be preserved under the harsh conditions of material shaping processes?

To answer this question, we have carried out a comprehensive investigation of the molecular and supramolecular states of acid-soluble collagen over a wide range of concentrations and temperatures. We draw, for the first time, a state diagram of collagen that encompasses the extreme concentrations and temperatures that characterize materials fabrication processes. Here, we explore the thermal events that determine the different state transitions of acid-soluble collagen, such as denaturation (on heating), crystallization and vitrification (on cooling), together with a concentration-dependent molecular to fibrillar transition. Using differential scanning calorimetry (DSC), rheological analysis, small- and wide-angle X-ray scattering (SAXS and WAXS, respectively) together with TEM, we have determined the characteristic temperatures of the system, as well as the supramolecular order of collagen molecules and fibrils.

The new constitutive state diagrams, both in solution and after fibrillogenesis, are novel tools to rationalize the design of collagen-based biomimetic materials, as illustrated in the last part of this article (see below). We anticipate that the fundamental insights drawn in this work will guide the processing strategies to develop materials with features that will approach, to a further extent, the outstanding properties of the ECM.

## Results and Discussion

### The state diagram of type I collagen in acidic solution

The state diagram of collagen in solution was studied over a wide range of concentrations, between 5.6 mg.mL^-1^ and 1350 mg.mL^-1^ (Fig. 1-A and B), and temperatures, between -80 and 80 °C. To investigate the collagen supramolecular arrangement as a function of the concentration samples at 40, 200, 400 and 900 mg. mL^-1^ were observed by PLOM between crossed polarizers. At 40 mg.mL^-1^ samples showed no birefringence due to the isotropic orientation of collagen molecules in solution at low concentration. The birefringence of samples at concentrations of 200 mg.mL^-1^ and above (Fig. S1) confirmed the presence of domains of aligned collagen molecules, in agreement with the widely described lyotropic liquid crystalline behavior of acid-soluble collagen^14,35,36^. Although the birefringence can be clearly observed above 200 mg.mL^-1^, the large sample thickness, around 200 µm, hindered the visualization of the fingerprint patterns characteristic of collagen mesomorphic phases. Phase contrast microscopy images (Fig. S1) confirm that the system is already heterogeneous at 200 mg.mL^-1^, with the detection of bundles that become increasingly intertwined and crowded at higher concentrations. These observations confirm the formation of complex aggregates of aligned collagen molecules, which are birefringent and have an optical refraction index different from that of the solution, as previously demonstrated for different acidic conditions^11^.

**Fig. 1.**
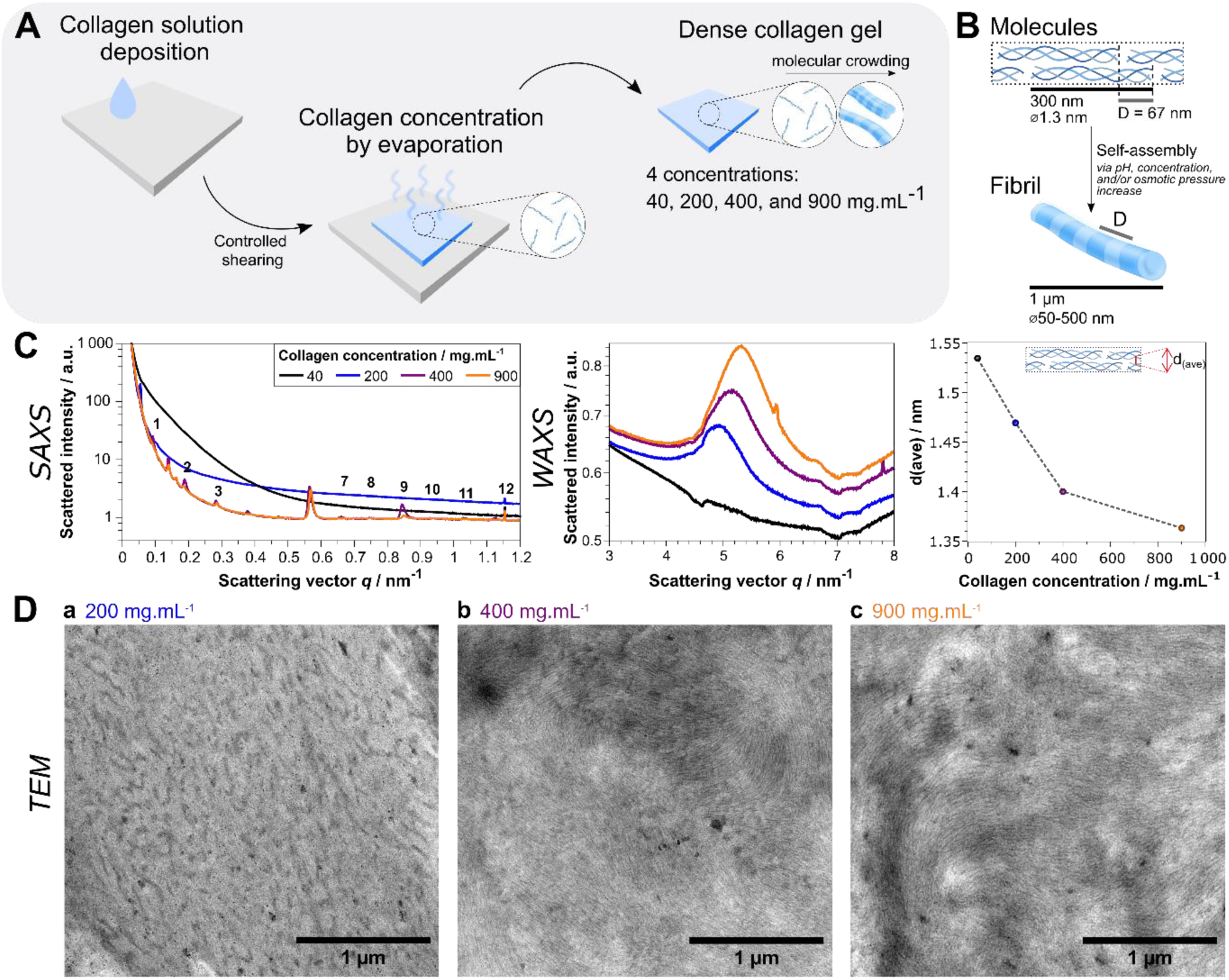
The supramolecular organization of type I collagen is determined by its concentration. Preparation and characterization of dense collagen samples at four concentrations: 40 (a-black), 200 (b-blue), 400 (c-purple), and 900 mg.mL^-1^ (d-orange). A) Thin films of dense collagen were prepared by drop casting, followed by shearing and controlled evaporation of the solvent. B) Scheme of the fibrillogenesis process *in vitro*. Characteristic dimensions were taken from Fratzl and Weinkamer^41^. C) Varying the concentration of collagen yields a range of different ultrastructures, characterized by small angle (left) and wide-angle X-ray scattering (middle). The dependence of the intrafibrillar lateral packing distance d(ave) (extracted from the WAXS data) on the concentration is shown on the right. (The artefacts observed at q = 7 nm^-1^ are due to the multichip nature of the detector.) D) TEM images of collagen in 3 mM HCl solution, as a function of the collagen concentration.

To probe the D-banding period and the intermolecular packing distance, we performed respectively SAXS and WAXS experiments over the whole collagen concentration range (Fig. 1-C). No diffraction peaks appear in the SAXS signal at 40 mg.mL^-1^ and only two weak peaks are observed at 200 mg.mL^-1^, demonstrating the lack of long-range order. A different behavior is observed at 400 and 900 mg.mL^-1^ where twelve equidistant diffraction peaks are detected, a signature of the 67 nm axial period found in native collagen fibrils^37^. Regardless of the dimensions and organization of the measured objects—fibril-like, pre-fibrils, or fibrils—these data demonstrate that 3D objects with a hierarchical organization are directly formed from collagen solution without pH or ionic strength variation. To our knowledge, this is the first report of a spontaneous transition from a molecular to a supramolecular organization of collagen molecules in hydrochloric acid solution^i^.

We probed the molecular packing between adjacent collagen molecules, and thus the compacity of the fibrillar units, using WAXS. As collagen concentration increased the peak shifted to higher scattering angles, indicating that the molecules become more tightly packed, reaching a final distance of 1.36 nm at 900 mg.mL^-1^. The molecular crowding resulting from increasing collagen concentration induces the formation of ordered molecular aggregates that reproduce the characteristic distances of native collagen fibrils, both in their D-banding pattern and in their intermolecular distances^10^. These changes at the molecular scale are accompanied by a sharp increase in the values of the elastic moduli, G’ and G’’, above 430 mg.mL^-1^ (Fig. S2). In addition to the characteristic distances deduced from the scattering experiments and the emergence of a gel-like behavior, ultrathin sections of 200, 400, and 900 mg.mL^-1^ samples were observed by TEM (Fig. 1-D). At 200 mg.mL^-1^, low-contrast objects are observed, with a diameter of 30 nm, suggesting the formation of supramolecular assemblies of collagen molecules. This observation confirms the WAXS data showing the lateral packing of collagen molecules. At higher concentrations (400 and 900 mg.mL^-1^), the observation of dense networks of nanofibrils confirms the characteristic 67 nm period measured by SAXS.

The results above, obtained at room temperature, were inserted in the temperature vs concentration state diagram of type I collagen in acidic solution (Fig. 2). In this diagram, we report the molecular organization of collagen as a function of concentration and temperature, both in a wide range. Contrary to the phase diagrams of collagen solutions reported to date—based on the characterization of the different mesophases as a function of acid concentration and ionic strength^10,38^—here we consider non-thermodynamic transitions like the molecular to fibrillar transition and vitrification, and other transformations, such as collagen denaturation. In solution collagen triple helices are thermally denatured when heated above ∼37 °C and between 50 °C and 60 °C for fibrillar collagen. Upon heating, the dissociation of the triple helices—assimilated to a melting process^39^—produces gelatin, which prevents the re-formation of the native fibrillar ultrastructure. Up to 100 mg.mL^-1^, the denaturation temperatures are close to 40 °C, corresponding to the denaturation of collagen in its molecular form^40^. For concentrations between 200 and 400 mg.mL^-1^, two denaturation processes were detected (Fig. S3-A). In this range, the upper denaturation temperature values are close to those found for collagen in its fibrillar form, confirming the existence of an ordered phase arising in solution, as inferred from the SAXS measurements. We hypothesize that at 200 and 400 mg.mL^-1^, the lowest denaturation temperature corresponds to the denaturation of collagen under its molecular form and the highest to an acidic fibrillar form. These observations lead to five distinct domains in the phase diagram of type I collagen in solution. The region above the denaturation temperature corresponds to the denatured form of collagen (D), i.e. gelatin. The three adjacent regions, below the denaturation line, correspond to molecular collagen (M), the coexistence of fibrillar collagen and denatured collagen (F+D), and a single acidic fibrillar collagen phase (F). The existence of a domain composed of F and D phases (between 40 and 60 °C), suggests that, below the denaturation temperature of molecular collagen, there is a domain where the molecular and fibrillar forms of collagen coexist. This domain, not previously reported, is bounded at high temperature by the denaturation of molecular collagen and at low temperature by the liquidus line in a concentration range between 200 and 400 mg.mL^-1^. A variety of materials processing techniques applied to collagen in solution result in a progressive increase of collagen concentration. Based on the different concentrations achieved during these fabrication processes, we are now able to determine the supramolecular state of collagen, opening a rational way do design biomimetic materials.

**Fig. 2.**
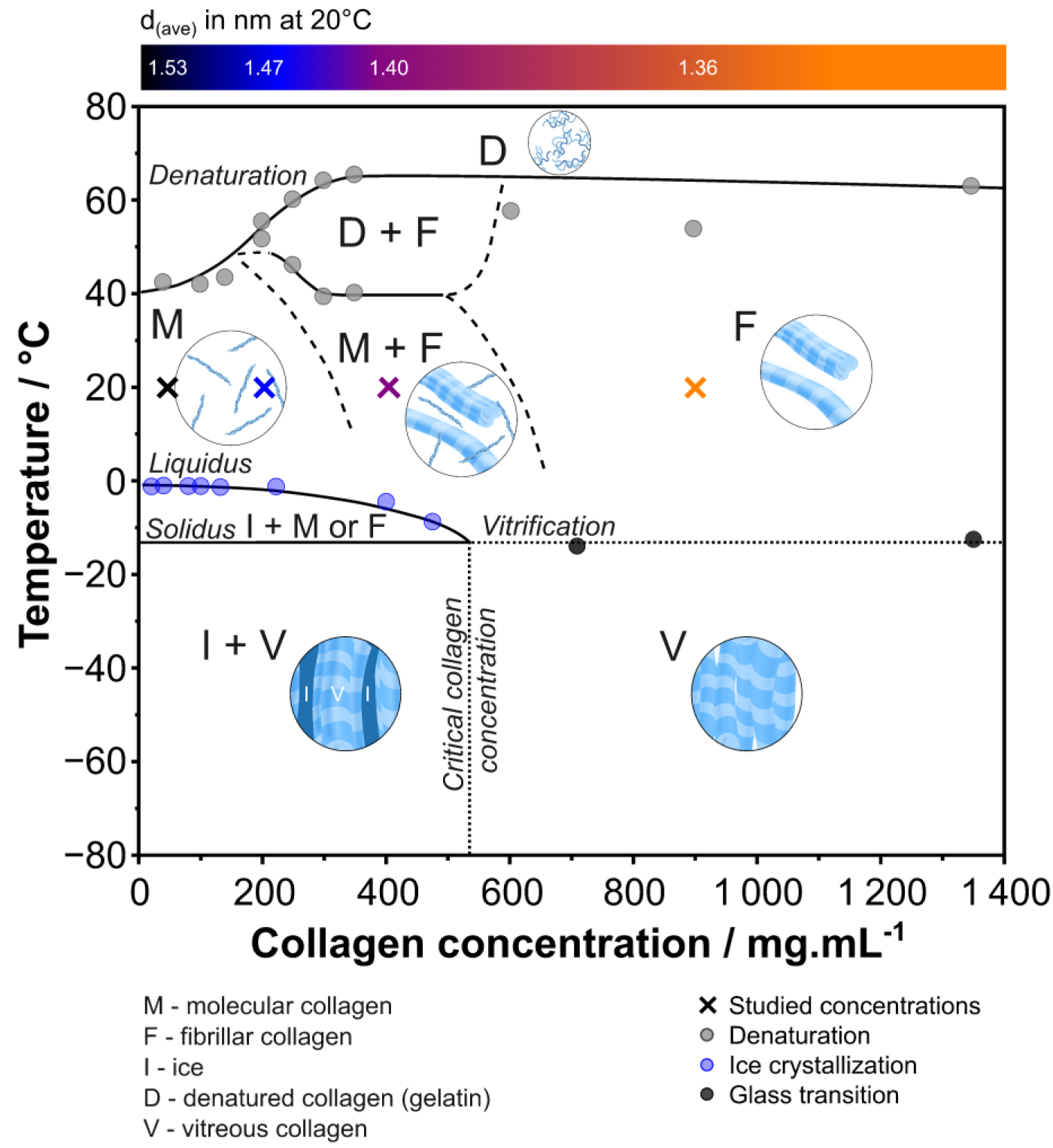
State diagram of acid-soluble type I collagen. The diagram combines data obtained from DSC, mDSC, SAXS, and WAXS. Details of the measurements used to describe the various state transitions are given in Fig. S3.

DSC experiments were also carried out to investigate the interactions of the solvent-solute system during freezing, a temperature regime that has been largely neglected in the literature. Solutions of different concentrations were frozen from 20 °C down to -80 °C to assess the crystallization temperature of free water (Fig. S3-B). When cooling down collagen samples, in a concentration range between 5.6 and 530 mg.mL^-1^, the formation of ice nuclei and their subsequent growth occurs at the solvent crystallization temperature lowered by the corresponding cryoscopic depression, defining the liquidus line. Since collagen, like most molecular and ionic species, is insoluble in ice, molecules are progressively expelled from the ice phase and concentrated in between the ice crystals at a critical collagen concentration. This concentration can be determined by DSC, as it results in the absence of an exothermic water crystallization peak, and was measured at 530 mg.mL^-1^ (Fig. S3-C). We consider this concentration as the point at which all remaining water is bound to collagen and cannot be further displaced to form ice crystals even at temperatures as low as -80°C. The absence of crystallization above this concentration suggests that between the denaturation temperature and -80°C the system is monophasic, composed of collagen and bound water.

We hypothesize that cooling collagen solutions above the critical concentration results in the formation of a vitreous state composed of collagen fibrils and disordered bound water. To verify this hypothesis, we measured the glass transition temperature (Tg) by modulated DSC (mDSC). The Tg for 704 mg.mL^-1^ was measured at *T*_*g*,_(704) = −13.9 °*C* (Fig. S3-D.b), a value remarkably close to that measured for dry collagen, at *T*_*g*,_(1350) = −12.5 °*C*, measured by dynamic mechanical analysis (DMA). Taken together, the data set obtained by SAXS, WAXS, DSC, mDSC and DMA lead, for the first time, to a complete picture of the different organization states of collagen over a wide range of concentrations (from 5.6 to 1350 mg.mL^-1^) and temperatures (from -80 to 80 °C). The state diagram provides a fundamental tool to describe collagen in solution, and it unveils the temperature and concentration-dependent self-assembly mechanisms that can prove instrumental in designing processing strategies to obtain materials that mimic the ECM.

### The state diagram of fibrillar collagen

Collagen fibrils are the central building block of native tissues. The previous results establish guidelines for handling collagen in solution, but they are insufficient to provide insights for collagen in its fibrillar form. Moreover, some materials processing techniques deal with previously fibrillated collagen materials, notably by compression^42,43^. To build a state diagram for fibrillar collagen, we immersed the collagen samples into PBS 10x for 24 hours to induce fibrillogenesis (Fig. 3-A). Observed under the TEM, negatively stained samples prepared at room temperature revealed fibrils with the typical D-banding pattern for all concentrations (Fig. 3-B). At low concentrations, small randomly dispersed fibrils are observed, whereas at 200 and 400 mg.mL^-1^ the resulting network is denser and composed of large striated fibrils. At 900 mg.mL^-1^, a tight network of nanofibrils forms, with barely visible striations. The reduced size of the fibrils is ascribed to the limited mobility of collagen molecules in solution at increasingly high concentrations, as confirmed by rheology measurements (Fig. S2). Such hindered mobility during fibrillogenesis results in the formation of smaller fibrils, featuring a smaller number of repeating units.

**Fig. 3.**
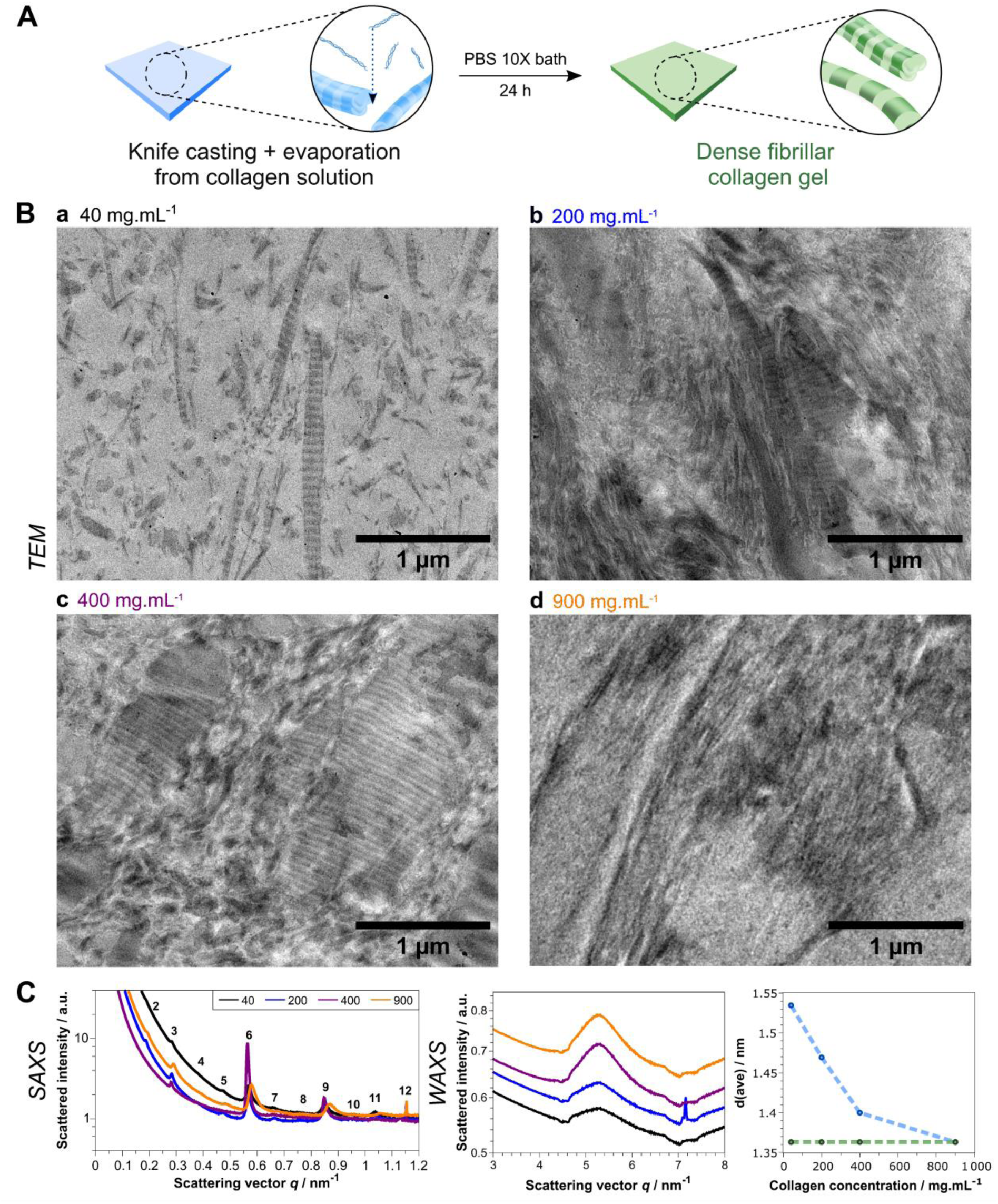
Elaboration and characterization of self-assembled dense collagen samples. A) Samples were immersed in a PBS 10x for 24 h. Four concentrations were studied: 40 (a-black), 200 (b-blue), 400 (c-purple), and 900 mg.mL^-1^(d-orange). B) TEM images of the samples displaying fibrils, and their arrangement as a function of collagen concentration. C) Left: Fibril formation was confirmed for all collagen concentrations by the native-like long-range order measured by SAXS. Middle: The molecular packing studied by WAXS is characterized by a constant average distance between fibrils, *d*_(*ave*)_ = 1.36 *nm*, independent of the initial concentration. Right: concentration dependence of *d*_(*ave*)_ for collagen in solution (blue points) and in fibrillar form (green points).

Observation of the samples by PLOM shows that, qualitatively, the birefringence increases with increasing concentration (Fig. S4). Moreover, samples at lower concentration (40 and 200 mg.mL^-1^) show a large contrast in light transmission depending on the angle between the substrate and the polarizer direction, indicating that the collagen is macroscopically aligned by surface effects. We hypothesize that, due to the higher packing and therefore limited mobility at high concentration, the chirality-induced twist of the collagen fibrils would be confined to the boundaries between domains, as suggested by de Sa Peixoto et al.^11^. This would lead to sharp twists in molecular alignment between domains, as found in plywood models.

At all concentrations investigated, the SAXS patterns of the samples show diffraction peaks corresponding to the characteristic D-banding of collagen fibrils. The positions of the diffraction peaks remain constant, but the peaks become sharper above 40 mg.mL^-1^ until 400 mg.mL^-1^, while at 900 mg.mL^-1^ the reflection width becomes larger. Altogether, the SAXS data correlate with the observations made by TEM. The relative intensities of the 67 nm harmonics depend on the concentration. In particular, at a concentration of 400 mg.mL^-1^, extinctions are observed for many peaks, such as the 4th and 5th orders, while the 6th order has an exceptionally high intensity. This peculiar behavior is reminiscent of previous observations of pathological fibrotic tissues (scars, burns or diabetic patients’ skin^44–46^, which show similar atypical patterns. In contrast to acid-soluble collagen, the average distance between the centers of the molecules, calculated from WAXS experiments at *d*_(*ave*)_ = 1.36 *nm*, is independent of the concentration. This observation suggests that fibrillogenesis results in a robust supramolecular assembly irrespective of the concentration. Interestingly, these values perfectly match the one found for collagen in solution at 900 mg.mL^-1^, proving that the crowding effect in solution suffices to assemble collagen molecules into tightly packed fibrils in the absence of a pH increase and at low osmolarity.

In parallel to the state diagram built for acid soluble type I collagen, we have built the state diagram of fibrillar collagen based on thermal analysis conducted by DSC and mDSC. The denaturation temperature, Td, of fibrillar collagen is approximately 55°C, which is similar to that of collagen in acidic solution at high concentrations. This similarity further supports the idea that a fibrillar state of collagen can be obtained from solution at high concentrations.

As for the state diagram of collagen at low temperature, we have followed the heat exchanges during freezing using DSC, which allows to determine the characteristic crystallization temperatures of collagen gels. The crystallization events, measured up to 400 mg.mL^-1^, were fitted with the modified Schroder-van Laar equation^47^. We estimated that the critical collagen concentration for fibrillar collagen should be somewhere above 400 mg.mL^-1^, the highest concentration at which water crystallization was observed. Above this critical concentration, fibrillar collagen solutions will turn directly into a vitreous state at a temperature Tg of -25 °C (Fig. S5-C). We have estimated the collagen critical concentration in its fibrillar form from the intersection of the Tg line and the fit of the liquidus data points, which should be close to [*col_gel_*]*_crit_* = 680 *mg*. *mL*^−1^, a value larger than that found for molecular collagen [*col_gel_*]*_crit_* = 530 *mg*. *mL*^−1^. Fibrillar collagen, due to the lateral association between adjacent collagen molecules, exposes fewer sites for interaction with bulk (or free) water molecules than its molecular counterpart. As a consequence, for the same analytical concentration, fibrillar collagen has a higher content of free water than collagen in solution. This explains why a larger quantity of free water can be frozen in fibrillar collagen gels than in the molecular counterpart.

Upon heating or cooling, all the characteristic temperatures of the state diagram are constant regardless of the collagen concentration, except for the liquidus curve, which depends on the cryoscopic depression induced by the solutes. The invariance of the temperatures with collagen concentration, in particular the denaturation temperature *T*_*d*_, indicates that the collagen fibrils have comparable thermal stability, irrespective of the concentration, contrary to what has been previously reported^48^.

These observations clarify the effects on the thermal stability of collagen fibrils of two variables that are usually intertwined: their size and their intrafibrillar molecular packing. The data collected here, in which the fibril size is variable (as observed by TEM) but the intermolecular packing is constant (as observed by WAXS), allow to pinpoint the primordial role of the molecular packing on the thermal stability of collagen fibrils. In addition to the fundamental aspects of collagen fibril stability discussed above, it seems clear that post-fibrillogenesis processing of collagen does not affect its molecular packing and thus the thermal stability of the resulting materials. This observation is particularly relevant to shaping processes such as compression (Fig. 4, green arrow)^49^.

**Fig. 4.**
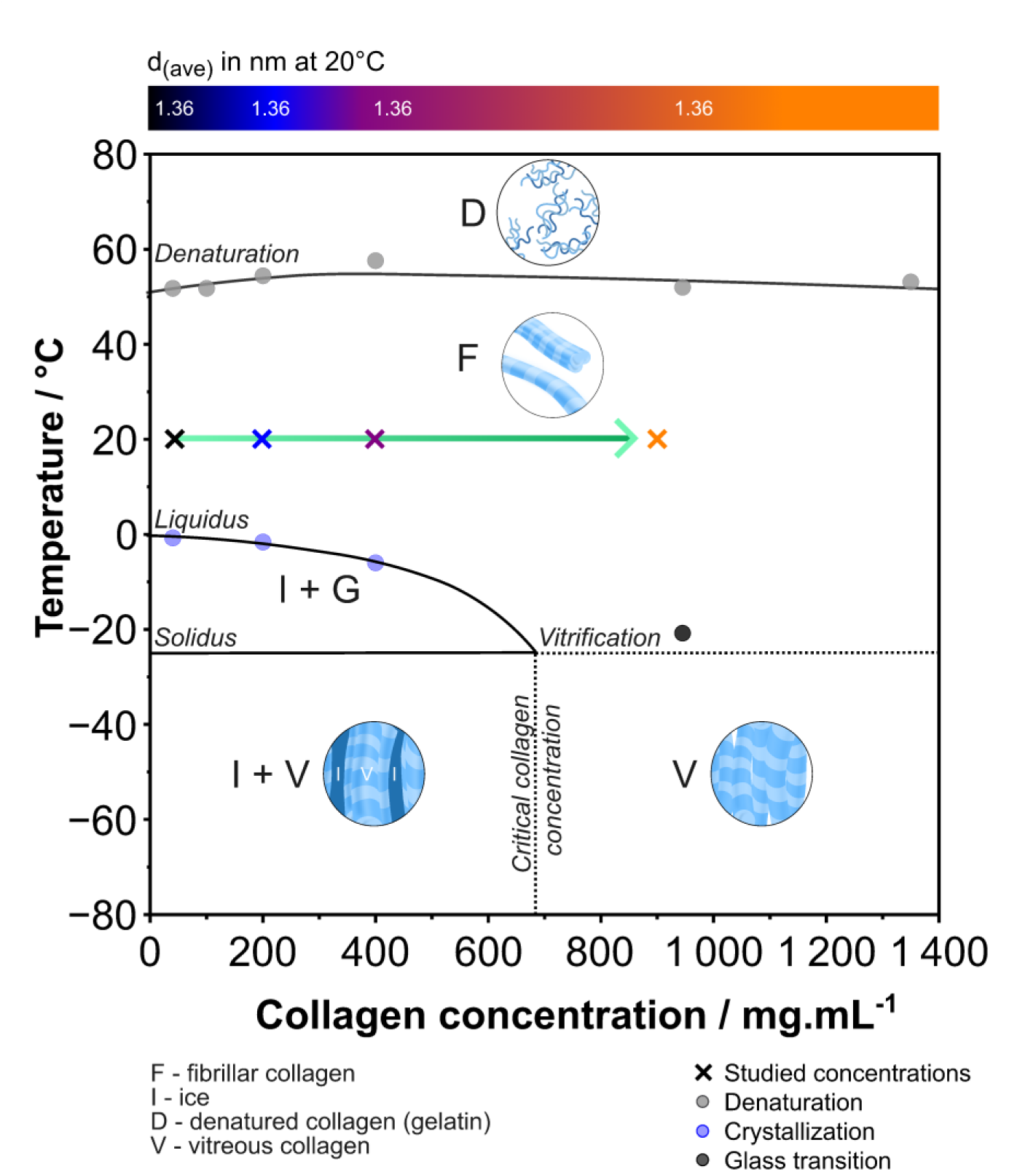
State diagram of fibrillar type I collagen. Dense collagen samples were fibrillated by immersion in 10x PBS. Details of the measurements used to describe the various state transitions are given in Fig. S5. Green arrow: path describing compression-based processing techniques^43^.

### Application of collagen state diagrams to the rational design of biomimetic materials

Taken together, the state diagrams described above define the conditions of temperature and concentration that control the molecular integrity and hierarchical organization of collagen, from the molecular packing distances up to the emergence of discrete fibrillar objects. This knowledge provides a new framework for building biomimetic materials that replicate the cellular micro-environment, both in terms of composition and functional properties. By tuning temperature and collagen concentration, and by considering the various state transitions mentioned above, the processes required to achieve the properties of native collagenous tissues can be rationalized. To illustrate this approach, we have used the state diagram of acid-soluble collagen to follow the events that occur during ice templating of a collagen solution at 40 mg. mL^-1^, in 3 mM HCl.

On cooling, from the first moments of ice formation, the local collagen concentration gradually increases along the liquidus line until it reaches the critical concentration of 530 mg.mL^-1^, at -12 °C. To visualize the events associated with this process, we have conducted controlled freezing of a collagen solution containing Rhodamine-B enclosed in a Hele-Shaw cell by sliding the sample from a hot plate at 20 °C to a cold plate at -30 °C under a cryoconfocal microscope^50^ (Fig. 5-C.a and Video S1). This series of events was plotted on the state diagram of collagen in solution to illustrate how the local composition of the collagen-rich fraction during freezing evolves (blue line in Fig. 5-A).

**Fig. 5.**
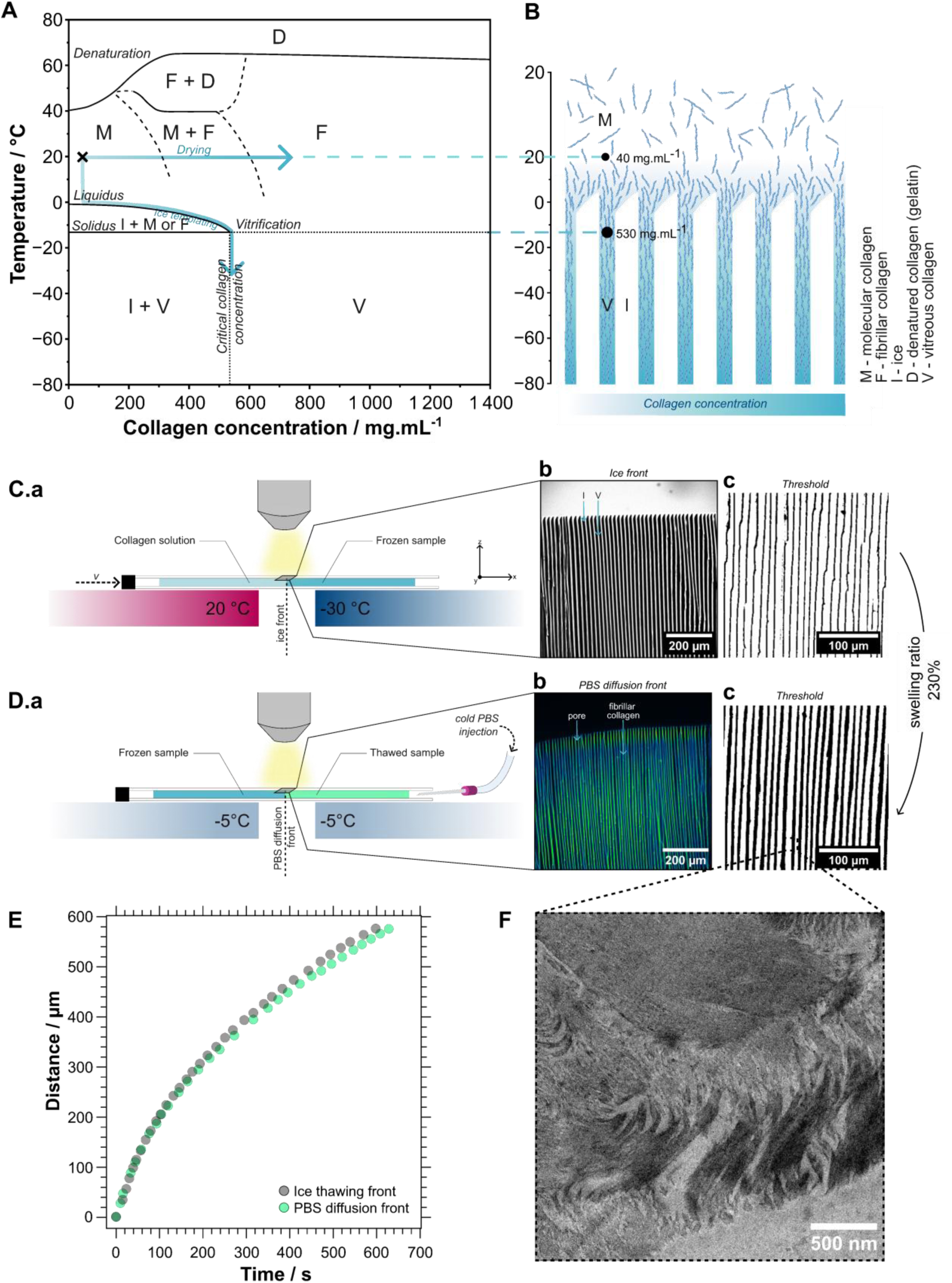
In situ cryoconfocal analysis of directional freezing and fibrillogenesis of a collagen sample at 40 mg.mL^-1^. A) Blue arrows track the fate of collagen in solution during drying (upper horizontal arrow) and ice templating (lower arrow). B) At 40 mg.mL^-1^ and 20 °C, collagen is in a molecular state. Below -1 °C, ice nucleates and grows into lamellar structures that expel collagen. In the interstitial space, collagen concentration increases, reaching the critical concentration of 530 mg.mL^-1^. C) a. The cryoconfocal set-up allows to freeze a solution by sliding samples between defined thermal boundary conditions (from 20 to -30 °C, 2 mm apart). b. Fluorescence image of a frozen sample showing the collagen solution (light gray) and the ice (black) fraction. c. Segmented image depicting the extent of phase separation. D) a. In situ topotactic fibrillogenesis of frozen collagen using PBS 10x solution, at a constant temperature of -5 °C, under the cryoconfocal microscope. b,c. Fluorescence and segmented image obtained after *in situ* fibrillogenesis. E) *In situ* measurement of the topotactic fibrillogenesis process allows to identify the kinetics of ice melting (1st front) and fibrillogenesis (E-2nd front). The TEM image of the fibrillated collagen wall (F) shows a dense network of small fibrils and demonstrates that the collagen can still self-assemble into fibrils by exposure to PBS 10x after ice templating.

After the freezing process we stabilized the collagen phase by topotactic fibrillogenesis^17^. Holding the sample at -5 °C after freezing allows the ice phase to remain stable until it comes in contact with the PBS solution. At this point, the pH change and the cryoscopic depression caused by the buffer triggers the fibrillogenesis of the collagen-rich fraction and the thawing of ice crystals, respectively. The relative kinetics of these events determines the ability of the process to stabilize the collagen walls and prevent their swelling. These two events were captured as image sequences (Video S2), allowing to measure their local velocity (Fig. 5-D.b,c). Fig. 5-E shows the progression of the ice thawing front (1st front) and the PBS diffusion front into the collagen walls (2nd front), as a function of time. Comparing these two fronts reveals that the ice thawing proceeds slightly faster than the PBS diffusion within the collagen walls. Although this difference seems modest, it has a significant impact on the interstitial zone, which undergoes partial swelling and redilution, bringing the local collagen concentration from 530 mg.mL^-1^ reached after freezing (Fig. 5-C,D) to 230 mg.mL^-1^ after fibrillogenesis.

The stabilization of collagen molecules into fibrils was confirmed by TEM (Fig. 5-F). The resulting gel shows a dense network composed of tightly packed fibrils of different sizes, reminiscent of the features observed at 200 mg.mL^-1^(Fig. 3-B.b). Larger bundles are found at the interface with the pores (bottom-right of the TEM image), where the re-dilution provides greater mobility for the molecules to self-assemble into larger supramolecular structures. A few cross-striated fibrils were also found across the walls. However, most assemblies are too small to generate the D-banding pattern, owing to the high density of collagen molecules. It is noteworthy that the harsh conditions of freezing do not alter the ability of collagen to form native-like fibrils when combined with appropriate fibrillogenesis conditions.

In addition to the ice templating process, the impact on the state of collagen can be assessed for other material processing strategies using the state diagrams described above. Electrospinning^22^, electroconcentration^31^, and spray drying^24,25^ are evaporation-based processes that have been successfully applied to the production of collagen materials at ambient temperature. In both cases, these transformations can be described as a horizontal line in the state diagram of collagen in solution, starting from the initial concentration (Fig. 5, blue arrow). We therefore anticipate that these processes have led to a sol-gel transition associated with tighter molecular packing during evaporation. Post-fibrillogenesis processes can also be interpreted as specific trajectories in the state diagram of fibrillar collagen. Among these, one of the most important is compression, where a mechanical constraint is applied to increase the concentration of fibrillar collagen gels to a desired value^43,49^ (Fig. 4, green arrow). Unlike solution-based processes, compression-based processes do not impose a significant change in the collagen organization in terms of molecular packing. The ability to interpret the effects of different processing strategies on the state of collagen using the diagrams described above provides an instrumental tool to guide the design of new collagen-based biomimetic materials.

### Conclusions

To the best of our knowledge, our study is the first to report the state diagrams of collagen in solution and in its fibrillar form in a particularly wide range of concentration and temperature.

In solution, at temperatures above 40 °C, collagen molecules denature into gelatin, preventing spontaneous self-assembly. Upon freezing, collagen segregates between ice crystals along the solidus line and gradually concentrates until a solubility limit, called the critical collagen concentration, at 530 mg.mL^-1^ is reached. While crossing this vertical line at low temperature, collagen reaches a vitreous state. At higher temperatures, above the solidus, increasing the concentration of collagen induces the formation of fibril-like structures. Scattering experiments, combined with microscopy techniques (PLOM, TEM), confirmed the self-assembly of molecules at concentrations ranging from 200 to 530 mg.mL^-1^.

Similarly, we built the state diagram of fibrillar collagen, induced by immersion in PBS 10x. For each concentration, the formation of cross-striated native-like fibrils was confirmed by both SAXS experiments and TEM imaging. In addition, immersion in PBS 10x results in a constant intrafibrillar packing distance across the entire concentration range. This fibrillogenesis process therefore leads to the formation of well-ordered supramolecular organizations, regardless of the initial concentration.

We anticipate that the state diagrams of collagen will provide guidance for the rational design of biomimetic materials built from the main protein of the ECM, ensuring that the features responsible for achieving the native ultra-structure of collagen are preserved. For example, ice templating provides a convenient way to segregate collagen between ice crystals, locally reaching very high concentrations. In addition, it allows for control over the material texture, which combined to topotactic fibrillogenesis ensures the formation of fibrillar collagenous materials, a step forward in mimicking organ structure and function. The state diagram of collagen in solution allowed to follow these events in detail and to describe the organization of the protein throughout the different stages of fabrication. The state diagrams obtained in this work also provide keys to rationalize fibrillar collagen compression processes. Finally, our approach can be extended to other biomolecules, including fibroin, fibrin/fibrinogen, chitin/chitosan and keratin. Such diagrams would improve our understanding of the conditions that govern their organization, and thus how close they are able to mimic the native tissues they derive from. We expect that such a predictive power will become instrumental in designing new materials processing strategies in the field of biomaterials.

## Acknowledgements

The authors kindly acknowledge C. Djédiat for the preparation of TEM samples, and SOLEIL for provision of synchrotron radiation facilities (under the approved proposal #20221057) and thank Thomas Bizien for assistance with the X-ray scattering experiments at the SWING beamline. IM acknowledges the PhD fellowship from the Physics and Chemistry of Materials’ graduate school (ED397). This work was supported by Agence Nationale de la Recherche (ANR) under grant agreement ANR-20-CE19-0029.

## Supporting information

### Experimental section

#### Fabrication of the collagen scaffolds

Type I collagen was extracted from the tendons of young rat tails according to a protocol adapted from that of Gobeaux et al.^51^. Tendons were thoroughly cleaned with phosphate buffered saline (PBS) 1X and 4 M NaCl solutions and dissolved in a 3 mM HCl solution. Differential precipitation with 300 mM NaCl and 600 mM NaCl, followed by redissolution and dialysis in 3 mM HCl, provided collagen of high purity. The final collagen concentration was determined using hydroxyproline titration and quantified at 5 mg.mL^-1^. The extracted collagen solution was then concentrated to 40 mg.mL^-1^ by centrifugation. The initial solution was transferred into Vivaspin tubes with 300 kDa filter and centrifuged at 3000g at 10 °C, until the final concentration was reached. Handling higher concentrations by centrifugation is technically challenging, since the increasing viscosity that follows collagen concentration hinders its use. To compensate this limitation, samples were concentrated by evaporation at ambient temperature, under sterile conditions, to reach concentrations of 200, 400, and 900 mg.mL^-1^, after deposition on a glass slide using a Doctor Blade® knife. The concentration after evaporation was calculated based on mass difference. The initial deposited volume was determined to reach a final weight of 50 mg for each concentration. As evaporation is faster at the sides of the sample, all characterizations in this study were performed near the center of the samples.

Two types of experimental conditions were compared: (a) collagen in solution, and (b) collagen fibrillated in PBS buffer. For (a), samples were simply stored in hermetically sealed petri dishes at 4 °C to minimize evaporation. Previous tests confirmed that no significant mass change was observed over time under these conditions. For (b), samples were immersed in a sterile PBS 10x buffer bath for 24 h. After rinsing with PBS 5x, gels were placed in PBS 5x buffer and stored at ambient temperature for 2 weeks to secure fibrillogenesis. At the end of the process, collagen gels were stored at 4 °C.

#### Small angle X-ray scattering (SAXS) and wide angle X-ray scattering (WAXS)

Customized square frames were printed in acrylonitrile butadiene styrene (ABS) filament using a Zortrax M-200 3D-printer to prepare cells for collagen samples during scattering experiments. First, a thin (ca. 23 µm) layer of Mylar was glued to one side of the frame, then the collagen solution or fibrillated sample was deposited on the Mylar layer. Finally, the cell was sealed by gluing a second layer of Mylar to the other side of the frame. Collagen solutions were concentrated directly in the (open) cell rather than on glass slides due to handling issues, as the gels were too sticky to be displaced, whereas fibrillar collagen samples were prepared aside and cut to fit in the frames.

To ensure the robustness of the measurements, X-ray scattering was recorded from 16 (4×4 grid) different points in each sample and three consecutive scattering images were recorded at each point. Since there was no difference between the first and last scattering pattern at each point, beam damage was ruled out.

SAXS and WAXS experiments were performed at the Swing beamline of SOLEIL, the French Synchrotron Radiation Facility (Saint-Aubin, France). The X-ray wavelength was λ = 0.1033 nm and the sample to detection distance was either 6.216 m (SAXS) or 0.518 m (WAXS). The scattered X-rays were collected with an Eiger-4M detector (pixel size of 150 µm) and the exposure time was typically 1 s. The X-ray scattering data was azimuthally averaged and normalized using the Foxtrot software developed at the beamline, resulting in plots of the scattered intensity versus scattering vector modulus, q (q = (4πsinθ)/λ where 2θ is the scattering angle) ranged between 2×10-2 and 8 nm^-1^. The background was subtracted after fitting by a polynomial function. Considering a hexagonal packing of the molecules, the average distance between the center of the molecules, d(ave) was derived from the peak position, qmax, in WAXS by the relation d(ave) ≈ 2π/qmax*(2/ 3).

#### Transmission electron microscopy (TEM)

Fibrillated and in solution samples were cross-linked with 2.5% paraformaldehyde (PFA), 2% glutaraldehyde, 0.18 M sucrose, and 0.1% picric acid for 12 h. The samples were then post-fixed with uranyl acetate in ethanol for 12 h, and dehydrated using baths of increasing concentrations in ethanol. Longitudinal sections of samples were embedded in SPURR-S resins prior to sectioning, transversal to the main axis. 70 nm ultrathin sections (Leica microtome) were contrasted with uranyl acetate and observed on a transmission electron microscope (FEI Tecnai Spirit G2) operating at 120 kV to observe the ultrastructural collagen features. Images were recorded with a CCD camera (Orius Gatan 832 ccdigital) at 6.5 kM.

#### Polarized light optical microscopy (PLOM) and phase contrast microscopy

Samples were embedded in 2.5% agarose and 200 µm semi-thin sections were cut with a vibratome (Compresstome). The sections were examined under two Olympus BX51 microscopes. For the PLOM, the microscope was equipped with an Olympus U-POC-2 (NA: 0.9) condenser and a 50X objective (Olympus LMPlan50XFL (NA 0.50)), and crossed polarizers to observe the birefringence of the sections. The phase contrast microscope was equipped with an Olympus U-PCD-2 (NA: 1.1) condenser and a 40X objective (Olympus PLCN40XPH (NA 0.65)).

#### In situ cryoconfocal microscopy

The confocal microscopy system designed to inspect the ice front during directional freezing^52^ consisted of a temperature controller connected to two independent Peltier elements (ET-127-10-13, Adaptive, purchased from RS Components, France) coupled with a controlled XY stage. The distance between the thermal elements was 2 mm. The freezing process was observed under a Confocal Laser Scanning Platform Leica TCS SP8 (Leica Microsystems SAS, Germany) using a 20x objective. For the analysis of the collagen wall width resulting from ice templating and the assessment of the fibrillogenesis impact on it, rhodamine B (0.05 mg per mg of protein) was dissolved in 4 wt% collagen solution. The resulting fluorescent solution was deposited on a dedicated Hele-Shaw cell (glass slide covered with a coverslip and sealed with a 100 µm-thick tape). The samples were moved from the ambient temperature to the cold Peltier elements (temperature range from 20 to -30 °C) by a stepper motor at 50 μm.s^-1^ linear velocities. The dimensions of the concentrated collagen walls were analyzed using FIJI software^53^. For fibrillogenesis imaging, the temperature was maintained at -5°C. Cold phosphate buffer solution (PBS) at a 10x concentration was injected with a needle in between the glass slide and the coverslip. The samples were then imaged during fibrillogenesis as the diffusion front progressed.

#### Differential scanning calorimetry (DSC)

The thermal transitions of collagen samples were measured by DSC. The samples (around 5 mg) were encapsulated in sealed aluminum pans and inserted in the chamber of a TA Instruments Q10 DSC under a dry nitrogen gas flow of 50 mL.min^-1^, coupled with a cooling flange. The instrument was calibrated using an empty cell baseline and an indium sample for heat flow and temperature. All measurements were performed at a cooling or heating rate of 5 °C.min^-1^. Denaturation temperatures (Td) were measured by heating from 10 to 180 °C. Crystallization temperatures (Tc) were measured by coupling cooling and heating cycles to minimize the supercooling effects occurring in aqueous solutions. A first cycle led to complete freezing of the free water, followed by heating up and cooling down until crystallization no longer occurred from a supercooled state, but from ice crystal seeds remaining from incomplete melting. The return point at which the equilibrium is in favor of the melting was defined as the Tc. The critical collagen concentration was determined by freezing increasing concentrations of collagen in solution, until no freezing event was observed. The glass transition temperature (Tg) was measured by heating solutions from -80 to 20 °C, using the temperature-modulated mode DSC (mDSC), and analyzing it in the reversing component. The experimental conditions (amplitude, frequency, and heating rate) were optimized and set at a temperature amplitude of 1 °C with a period of 60 s.

#### Establishing the liquidus lines

In solution the solvent crystallization temperatures decrease from -0.6 °C at 5.6 mg.mL^-1^ to -8.7 °C at 475 mg.mL^-1^, due to the cryoscopic depression of the collagen solution, which allows the positioning of the liquidus line of the diagram, a transition line from one phase (collagen solution, S) to two phases (ice and concentrated collagen solution, I+S). This transition can be described theoretically by a modified Schroder-van Laar equationw^47^ (eq. S1), established for synthetic polymers, which was fitted to the experimental data:

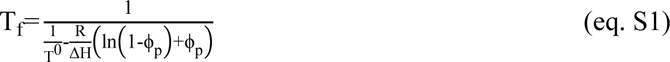

where T_f_ is the freezing temperature in Kelvin, ϕ_p_ is the collagen volume fraction (Table S1), T^0^ is the solvent freezing temperature, R is the molar gas constant, ΔH is the solvent melting enthalpy (ΔH(3 mM HCl) = 5752.2 J.mol^-1^), and N is the degree of polymerization of the polymer, which was considered here as N=∞.

#### Dynamic mechanical analysis (DMA)

The measurement of the Tg of dry collagen, was measured by DMA instead of DSC, since this technique could not detect the limited heat flow associated with this transition. Small-amplitude linear oscillations were applied to measure the dynamic moduli E, E’ and tanδ of a dry collagen solution brought to its solubility limit, on a DMA 850 from TA Instruments. A 3-point bending clamp of 17.5 mm was used. The strain rate was fixed at 2% per minute, with a frequency from 1 to 10 Hz. Measurements were repeated 3 times to confirm the glass transition temperature value (data not shown).

#### Rheology

Shear-strain oscillatory measurements were performed on collagen solutions with an Anton Paar rheometer MCR-302. An 8 mm plane-plane geometry (PP08/S) was fitted with a rough surface to prevent gel slippage. All measurements were performed at ambient temperature. The storage modulus G’ and loss modulus G’’ were recorded during a frequency sweep from 0.1 to 10 Hz with an imposed strain of 1%. The gap was set to ensure a minimal normal force of 0.01 N. Three samples of each solution were tested and averaged.

### Limitations of the study

This study is the first to establish the collagen state diagrams as a function of concentration and temperature, both in solution and in the fibrillated state. However, several limitations must be taken into account:

i. Accuracy of the determination of the concentration after evaporation. The concentration of the samples after solvent evaporation in ambient air was simply quantified by weighing. The concentration homogeneity of the samples at 40 and 900 mg.mL^-1^ is reliable, since in the the first case the sample is not dried at all and, in the second case, the sample is completely dry, as no further evaporation can take place in air. However, for the intermediate concentrations of 200 and 400 mg.mL^-1^, the final concentrations are only average values that were determined by ignoring any potential concentration gradient at the sample edges resulting from the partial drying process. To limit concentration errors, measurements were systematically made at the center of the samples (i.e. in the least evaporated part).
ii. State diagram boundaries. The intersection of the fitted liquidus line with the Tg line is an approximation. For acid-soluble collagen, the line describing the transition from molecular to fibrillar state has only been determined from ambient temperature experiments and is therefore also approximate. More experimental data at different temperatures would be needed to determine this transition precisely, in particular using in-situ SAXS measurements during freezing. State diagrams are drawn for a given concentration and type of acid. Varying these parameters can modulate the self-assembly process11 and shift the phase transitions. The change in position of the boundaries can also occur due to changes in heating and cooling rates, especially for the Tg line which is kinetics dependent. As they depend on the Tg line, the dotted lines are also kinetic-dependent transitions and therefore their positions are likely to shift under different temperature rate analysis conditions.
iii. Determination of the denaturation temperature. Due to the coexistence of two states in solution and the overlap of the denaturation temperature peaks, the Td were measured at the peak maxima rather than at the onset, as in classical measurements.
iv. Three-state intersection. The intersection of the regions of denatured fibrils (Td line), fibrillar collagen and denatured collagen (F+D), and fibrillar collagen (F), is not precisely known due to the lack of measured Td points and the difficulty in resolving thermal transitions with small temperature differences by DSC. Care must be taken in this region regarding the exact state of collagen.
v. Liquid-crystalline phases. In this work, we have focused on the state transitions of collagen rather than those involving liquid-crystalline phases. These have already been extensively described in terms of collagen and acid concentrations9. Extending their study over a range of temperatures would require equilibrium conditions, unlike material processing techniques which are mostly carried out under out-of-equilibrium conditions. An assessment of the liquid crystal-like organization of collagen as a function of temperature was therefore simply not within the scope of this study.

## Supplementary Figures

**Fig. S1.**
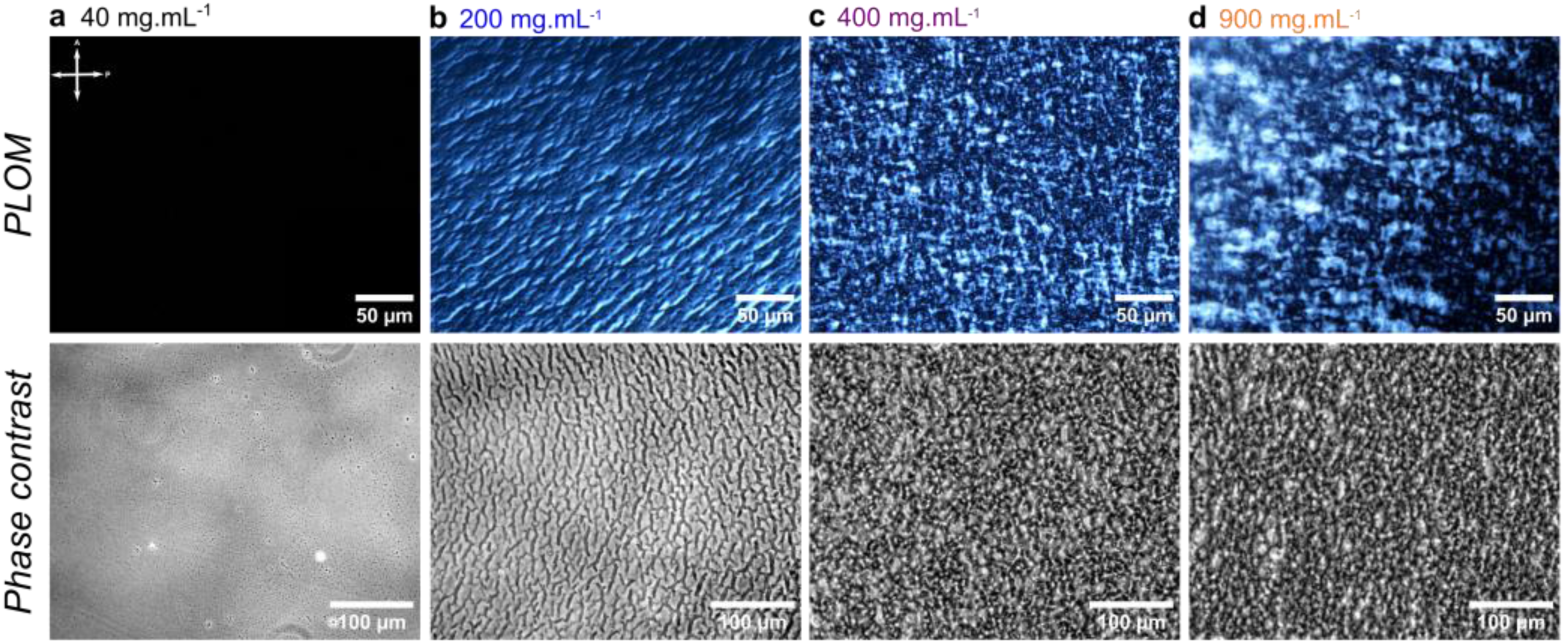
Observations of dense collagen samples from acid-soluble type I collagen at four concentrations: 40 (a), 200 (b), 400 (c), and 900 mg.mL^-1^(d). Increasing the collagen concentration leads to the appearance of birefringence and an increased complexity in the patterns observed by polarized-light optical microscopy (PLOM) and phase contrast optical microscopy.

**Fig. S2.**
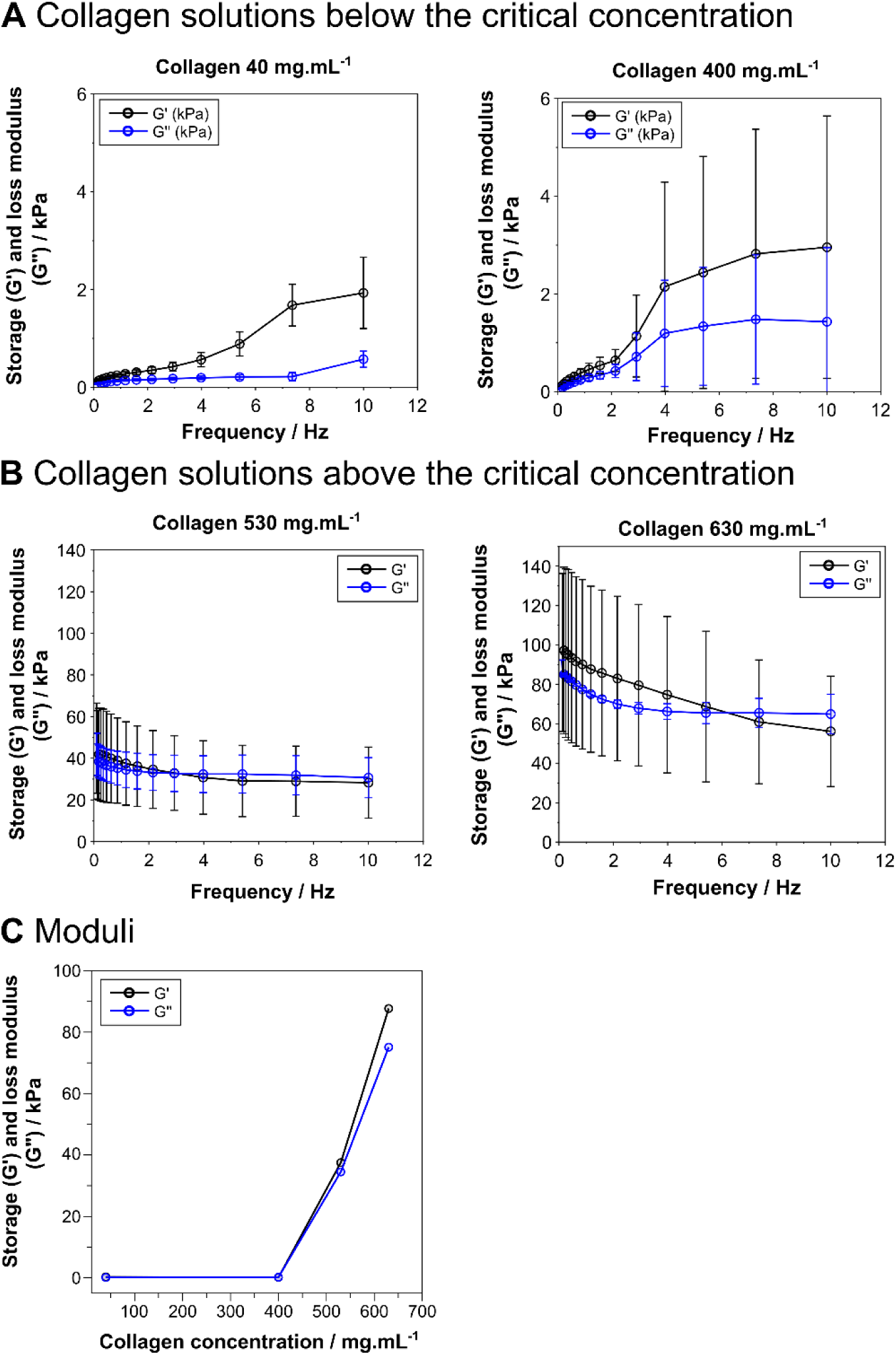
Variation of storage modulus G’ and loss modulus G’’ with frequency and collagen concentration for solutions below (A) and above (B) the critical concentration of 530 mg.mL^-1^. The G’/G’’ ratio, calculated for a frequency of Ꞷ = 7.36 Hz, shows a change in the rheological behavior associated with the sol-gel transition.

**Fig. S3.**
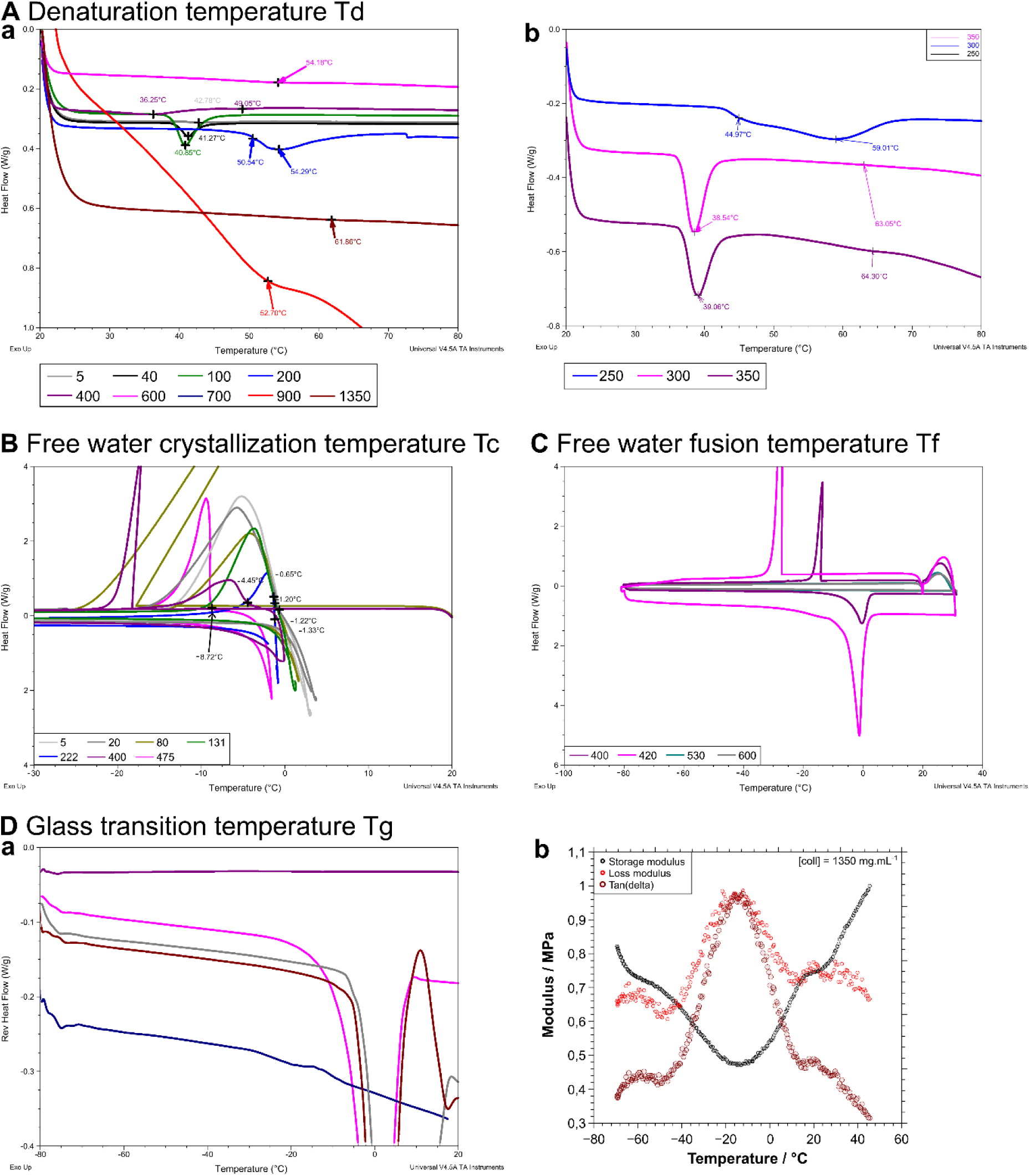
Analysis of the thermodynamic phase boundaries by DMA, DSC, and mDSC of collagen in solution. All measurements were performed at a rate of 5 °C.min^-1^. (A) Denaturation temperature Td, a) full range of concentrations, b) detail for 250, 300 and 350 mg.mL^-1^ (B) free water crystallization temperature Tc, (C) free water melting temperature measurements used to determine the critical collagen concentration, and (D) glass transition temperature Tg measured by mDSC (D-a) and DMA (D-b).

**Table S1.** Collagen volume fraction ϕp used in the modified Schroder-van Laar equation to fit crystallization temperature values.

| | Collagen concentration / mg.mL<sup>-1</sup> | Collagen volume fraction $\phi_p$ | T / K | T / °C |
| --- | --- | --- | --- | --- |
| Collagen in acidic solution | 0 | 0.000 | 273.15 | 0 |
|  | 5.6 | 0.004 | 272.54 | -0.61 |
|  | 20.5 | 0.015 | 271.95 | -1.20 |
|  | 40 | 0.030 | 272.14 | -1.01 |
|  | 80 | 0.059 | 272.04 | -1.11 |
|  | 100 | 0.074 | 272.03 | -1.12 |
|  | 131.7 | 0.098 | 271.82 | -1.33 |
|  | 222 | 0.164 | 271.93 | -1.22 |
|  | 400 | 0.296 | 268.7 | -4.45 |
|  | 475 | 0.352 | 264.43 | -8.72 |
| Collagen fibrillated in PBS 10x | 0 | 0.000 | 273.15 | 0 |
|  | 40 | 0.030 | 272.42 | -0.73 |
|  | 200 | 0.148 | 271.55 | -1.60 |
|  | 400 | 0.296 | 267.17 | -5.98 |

**Fig. S4.**
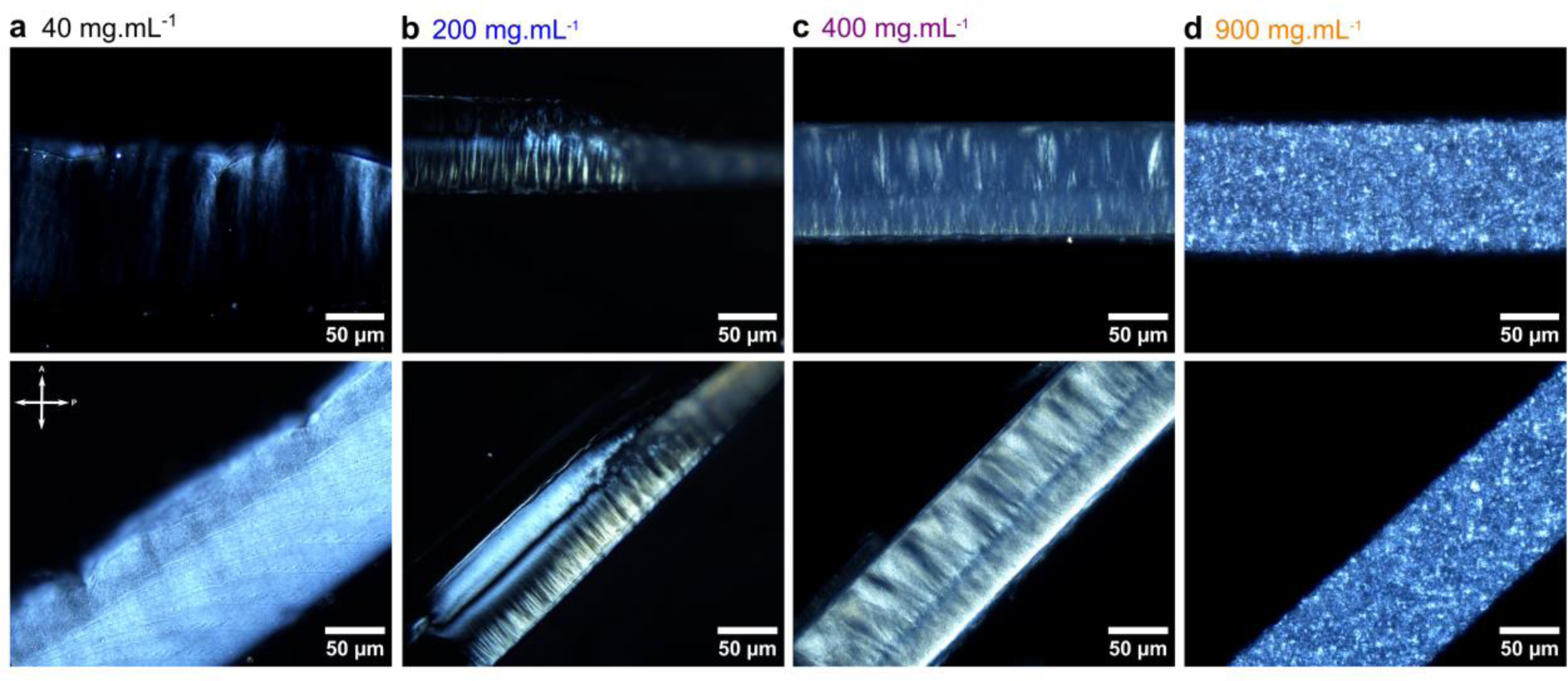
Polarized light optical microscopy images of transverse cross-sections of dense collagen samples fibrillated in 10x phosphate buffer solution at different concentrations. The sample surface was set parallel (resp. at 45°) to the polarizer direction in the top (resp. bottom) row. (In the top row, the free surface of the samples is at the top.)

**Fig. S5.**
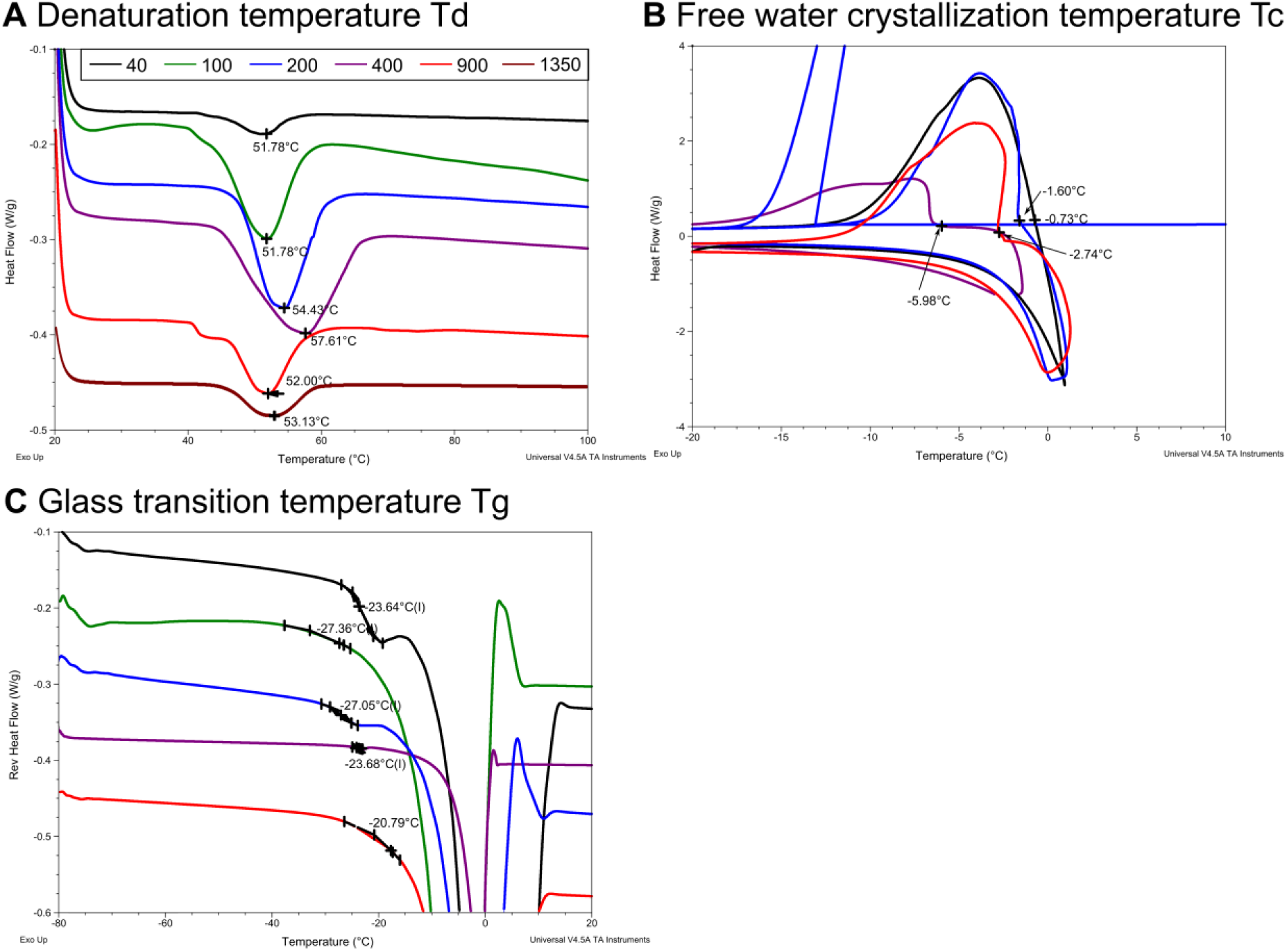
Analysis of the thermodynamic phase boundaries by DSC and mDSC of collagen samples fibrillated by immersion in phosphate buffer solution. All measurements were performed at a rate of 5°C.min^-1^. (A) Denaturation temperature Td, (B) free water crystallization temperature Tc, and (C) free water melting temperature measurements used to determine the critical collagen concentration.

**Video S1.** Sequential 2D fluorescence imaging of the freezing front of a 40 mg.mL^-1^ collagen solution frozen at 50 m.s^-1^. The collagen solution (light gray) is segregated in the interstices between ice crystals (black).

**Video S2.** Sequential 2D fluorescence imaging of the fibrillogenesis medium diffusion front in the previously ice-templated collagen solution, at a constant temperature of -5 °C. The first front corresponds to the melting of ice (black), which occurs before the self-assembly of the collagen molecules in the wall (green-blue).

## Footnotes

i However, SAXS experiments conducted in acetic acid at a concentration of 500 mM, have revealed the presence of the characteristic 67 nm period at 145 mg.mL-1. The shift in the concentration at which collagen assembles to form supramolecular objects is known to be modified by the type of acid and its concentration^10^.

